# Characterization and Evaluation of Hypoxia and LPS Preconditioned Placental Stem Cell Extracellular Vesicles in Immune and Intestinal Cells

**DOI:** 10.64898/2026.11.12.766307

**Authors:** Anchala Guglani, Maryssa A. Ellison, Victoria G. Weis

## Abstract

**Background:** Human placental stem cells (hPSCs) have demonstrated therapeutic potential in experimental necrotizing enterocolitis (NEC). Extracellular vesicles (EVs) secreted by the hPSCs contribute to these effects through paracrine signaling, however the molecular composition and biological activity of EVs can be influenced by culture conditions of the hPSCs. This study investigated whether hypoxic or lipopolysaccharide (LPS) preconditioning of the hPSCs alters the characteristics, protein composition, and biological activity of hPSC-derived EVs.

**Methods:** hPSCs were cultured under normoxic conditions, exposed to hypoxia (5% O_2_), or preconditioned with LPS (50 µg/mL). EVs were isolated from conditioned medium using ultrafiltration and size-exclusion chromatography, followed by characterized via nanoparticle tracking analysis, transmission electron microscopy, and single-particle detection of CD9, CD63, and CD81. Biological activity was evaluated in HIEC-6 intestinal epithelial cells through MTT, scratch wound closure, and nitrite detection assays. Immunomodulatory responses were examined in primary healthy neonatal rat CD4^+^ T-cell cultures using flow cytometry.

**Results:** EV preparations showed comparable particle concentrations and size distributions across all three culture conditions, with no statistically significant differences. Vesicle-like morphology and EV-associated tetraspanins were detected in all preparations. All three EV groups improved epithelial metabolic activity following LPS exposure, enhanced scratch wound closure, and reduced nitrite accumulation compared with their respective controls. Flow cytometric analysis showed no statistically significant differences in TNF-α, IL-17A, or FoxP3 expression among EV-treated CD4^+^ T-cell groups.

**Conclusions:** Hypoxic and LPS preconditioning did not significantly affect the measured physical characteristics or secretion concentrations of the EVs. Although EVs from all three conditions showed beneficial effects on intestinal epithelial cells, neither preconditioning approach demonstrated a clear functional advantage over naïve EVs in these simplified culture model. These findings suggest that preconditioning influences EV molecular cargo but more comprehensive biological function assays are required to delineate potential differences in effects on intestinal epithelial or T-cell functions. These data provide a basis for further studies examining its potential relevance to EV-based therapies for NEC.

## Introduction

Necrotizing enterocolitis (NEC) is a devastating inflammatory intestinal disease that primarily affects preterm infants. Immaturity of the intestinal epithelial barrier and mucosal defenses increase susceptibility to injury. Translocation of luminal bacteria across an injured barrier can then trigger a damaging local inflammatory response. Despite advances in neonatal care, treatment options for NEC remain limited and primarily focus on managing complications rather than addressing the underlying intestinal injury^1, 2^. This has led to growing interest in regenerative therapies that can reduce inflammation, preserve intestinal epithelial integrity, and promote tissue repair^1, 3^.

Paracrine signaling is considered an important mechanism underlying the therapeutic effects of stem cell-based and cell-free approaches in NEC^3^. This has led to growing interest in the stem cell secretome, which consists of soluble factors and extracellular vesicles (EVs) released by cells, as a potential cell-free therapeutic approach. Previous preclinical studies have shown that stem-cell-derived EVs can reduce intestinal injury and support epithelial repair in experimental NEC^4, 5^. In a neonatal rat model of NEC, administration of human placental-derived stem cells improved intestinal morphology and supported recovery of the intestinal epithelium and stem-cell niche^1^. These findings highlight the regenerative potential of placental stem cells. While this repair was induced via paracrine signaling, the specific secreted factors responsible for their beneficial effects remain unclear. One potential paracrine signaling contributor is EVs. EVs are membrane-enclosed particles that carry bioactive molecules, including proteins, lipids, and nucleic acids, which can influence the function of recipient cells^6, 7^. Supporting their potential role in intestinal repair, EVs derived from human amniotic fluid stem cells have been shown to reduce intestinal injury and inflammation while increasing markers of epithelial proliferation and intestinal stem cells in experimental NEC^7^. Similarly, EVs derived from therapeutic-grade placental stromal cells have been reported to regulate immune-cell proliferation and promote angiogenic activity in vitro, with proteomic analysis identifying proteins associated with immune signaling and cell migration^4, 8^. Together, these findings suggest that placental-cell-derived EVs may contribute to tissue repair and immune regulation, although their biological activity may vary depending on the cell source and recipient-cell environment^9^.

The properties of stem-cell-derived EVs are influenced not only by their cellular origin but also by the environment in which the parent cells are cultured. Modifying culture conditions through preconditioning may therefore provide a way to tailor EV composition and biological activity to a specific injury or disease setting. Hypoxia and lipopolysaccharide (LPS) are particularly relevant in NEC research, as both are commonly used in experimental models to reproduce aspects of intestinal injury and inflammation^1^. Previous studies have shown that hypoxic conditioning of human induced pluripotent stem cells altered the angiogenic activity of their EVs without significantly affecting EV production across different oxygen conditions^10^. Similarly, EVs derived from LPS-pretreated periodontal ligament stem cells showed different effects on inflammatory responses, epithelial barrier function, and cellular signaling compared with EVs from untreated cells in an intestinal inflammation model^11^. More recently, thrombin-preconditioned mesenchymal stromal-cell EVs demonstrated differences in proteomic profiles and biological activity compared with unconditioned EVs in experimental NEC^4^. Together, these studies demonstrate that preconditioning influences EV cargo and function, although its effects may vary with the conditioning stimulus, cell source, and disease application. Whether hypoxia or LPS preconditioning produces similar changes in EVs derived from human placental stem cells remains unclear. In our previous in vivo studies, EVs derived from human placental stem cells reduced epithelial injury, increased active and reserve intestinal stem-cell populations in a neonatal rat model of NEC^1^. These findings provide a basis for further investigating placental stem-cell-derived EVs and determining whether different preconditioning conditions alter their molecular composition and biological activity.

In the present study, we compared EVs collected from human placental stem-cell cultures maintained under normoxic conditions with EVs collected from cultures exposed to hypoxia or LPS. We characterized their particle concentration, size distribution, and morphology. We further evaluated the effects of these EVs on intestinal epithelial cells (HIEC-6) using cell viability, scratch wound healing, and nitrite detection assays. Their effects on immune-cell responses were examined using CD4^+^ T-cells enriched from healthy neonatal rat spleens, followed by EV treatment and flow cytometric analysis. Together, these analyses aimed to determine whether hypoxic and inflammatory preconditioning produces EV preparations with distinct molecular profiles and biological activities that may inform the development of placental stem-cell-derived EV therapies for NEC.

## Methods

### Human Placental Stem-Cell Isolation, Culture and Conditioning

Under an approved protocol by the Institutional Review Board of Wake Forest University School of Medicine, consent was obtained from patients prior to delivery for the collection and use of full-term human placenta specimens. Following placental tissue collection, processing was performed by the Manufacturing Development Center (MDC) within the Wake Forest Institute for Regenerative Medicine (WFIRM), Wake Forest University School of Medicine. As previously described^1^, chorionic tissue was digested for the isolation of perinatal placental cells and stocks of human placental stem cells (hPSC) were cryopreserved for subsequent use following expansion and c-kit positive selection. Cells were thawed and plated in tissue culture flasks with α-MEM media (HyClone, Logan, UT), 10% fetal bovine serum (FBS), and 1× Penicillin-Streptomycin. Cells were cultured at 37°C, 5% CO2, and no more than 80% confluency. Hypoxia-conditioned cultures were maintained at 5% O2 using a BioSpherix Xvivo system. LPS-conditioned cultures were exposed to LPS (50 µg/mL; L3012, Sigma, MO). Media was exchanged for media containing extracellular vesicle-depleted FBS and incubated for 48 hours. The 48-hour conditioned media (CM) was collected and centrifugated at 300×g for 10 minutes to remove cell debris. CM supernatant was collected to not disturb any cell debris pellet, transferred to sterile polystyrene bottles, and placed in -20°C until further processing (less than 6-months storage).

### Human Placental Stem-Cell Extracellular Vesicle Isolation and Characterization

CM was further processed for hPSC extracellular vesicle (EV) isolation by ultrafiltration using Amicon Stirred Cell 100kD ultrafiltration discs (MilliporeSigma, Burlington, MA), and concentration using Amicon Ultra-15 Centrifugal Filter Units (MilliporeSigma, Burlington, MA). Following filtration and concentration, 500 µl of condensed CM was processed via size exclusion chromatography using qEV2 / 35 nm Gen 2 columns (Izon Science, Medford, MA) to achieve high concentrations of small EV isolations within a recovery range of 35-350 nm.

Nanoparticle tracking analysis (NTA) (NanoSight Pro, Malvern Panalytical, Westborough, MA) was utilized to determine the size and concentrations of hPSC-EV isolations using the following parameter settings: blue 405 laser in scatter mode, camera level 12 (NTA 3.0 levels), shutter of 600, slider gain of 350, 25 frames per second at a total of 1499 frames captured, detection threshold of 5, and max jump distance at auto 12.9-13.9 pixels. Presence of transmembrane tetraspanin proteins (CD9, CD81, CD63) was confirmed using the ONI EV Profiler 2 Application Kit (following manufacturer’s instructions) with stochastic optical reconstruction microscopy on a Nanoimager (ONI, San Diego, CA). Transmission electron microscopy (TEM) confirmed EVs within the desired size range of <300nm and expected EV morphology. These EV characterizations of size, morphology, concentration, and tetraspanin expression are in accordance with MISEV2018 guidelines described by the International Society for Extracellular Vesicle^12^.

### *In Vitro* Assessment of EVs Bioactivity in HIEC-6 Cells

#### HIEC-6 Cell Culture and Cell Viability Assay

Human intestinal epithelial cells (HIEC-6; ATCC CRL-3266) were cultured in OptiMEM Reduced Serum Medium (Gibco Catalog No. 31985) based medium supplemented with 10% FBS, 20 mM HEPES, 10 mM GlutaMAX, 10 ng/mL epidermal growth factor, and 1% penicillin–streptomycin. Cells were maintained at 37 °C in a humidified incubator with 5% CO_2_. Cells at passage 21 were used for subsequent *in vitro* experiments.

HIEC-6 cells were seeded in 96-well plates and cultured until approximately 70% confluence. Cells were treated with increasing concentrations of lipopolysaccharide (LPS; 0, 2, 4, 6, 8, and 10 µg/mL; Sigma-Aldrich, Cat. No. L3012) for 24 hours across four treatment groups: LPS alone, LPS with EVs derived from naïve, LPS with EVs derived from LPS-preconditioned, and LPS with EVs derived from hypoxia-preconditioned. EVs were administered at 7.5 × 10^8^ particles/well. After treatment, 10 µL of MTT solution (5 mg/mL) was added to each well containing 100 µL of fresh medium and incubated at 37°C for 2 hours. Formazan crystals were solubilized using 10% SDS in 0.01 M HCl, and absorbance was measured at 570 nm using a microplate reader. Cell viability was expressed as a percentage of the untreated control^13^.

#### Scratch Wound Healing Assay

HIEC-6 cells were seeded in 96-well plates and cultured until confluent. A straight scratch (wound) in the center of each well was created using a sterile P10 pipette tip. Cells were treated with naïve, LPS-preconditioned, or hypoxia-preconditioned EVs (7.5 × 10^8^ particles/well). Untreated cells served as controls. Wound closure was monitored at 0, 12, 24, and 36 hours using an EVOS imaging system. Images were analyzed using ImageJ software by measuring scratch width at five locations per well. The mean scratch width was used to calculate the percentage of wound closure relative to the initial measurement (0 hour).

#### Griess Assay for Nitrite Detection

HIEC-6 cells were seeded in 96-well plates and incubated overnight at 37°C with 5% CO_2_. The culture medium was replaced with serum-free medium, and cells were treated with LPS (2 or 4 µg/mL) along with naïve, LPS-preconditioned, or hypoxia-preconditioned EVs for 24 hours. Untreated cells served as controls. Following the treatment, 50 µL of culture media supernatant was collected and mixed with 20 µL of Griess reagent. Samples were incubated for 30 minutes at room temperature in the dark, and absorbance was measured at 540 nm using a microplate reader. Nitrite levels were expressed relative to the corresponding LPS-only control, set at 100%.

### Neonatal Rat CD4+ T-Cell Isolation, EV Treatment, and Flow Cytometric Analysis

Spleens were harvested from 4-day-old healthy breastfed neonatal rats and processed into single-cell suspension using EasySep™ buffer (STEMCELL Technologies, Cat# 20144). Cell suspensions were filtered through a 70 µm strainer and centrifuged at 300 × g for 10 min to collect lymphocytes pellet. The cell pellet was resuspended, and CD4+ T-cells were enriched by negative selection using the EasySep™ Rat CD4+ T-Cell Isolation Kit (STEMCELL Technologies, Cat. No. 19642). Following negative selection, enriched cells were cultured in TexMACS™ (Miltenyi Biotec, Cat# 130-097-196), medium supplemented with 1X antibiotic–antimycotic solution (ThermoFisher Scientific), 1X 2-mercaptoethanol (ThermoFisher Scientific), and 5 U/mL recombinant human IL-2 (ThermoFisher Scientific, Cat# 200-02-10UG). Cells were seeded in 96-well U-bottom plates and activated with CD3/CD28 Dynabeads (ThermoFisher Scientific, Cat# 11456D) at a bead-to-cell ratio of 1:5. Activated and non-activated cultures received a single dose of hPSC-derived EVs (1,000 particles/cell) or saline control and were incubated for 48 hours at 37°C.

For flow cytometry, the cell cultures were incubated with GolgiStop™/GolgiPlug™ for 4 hours prior to collection. Following viability staining (BD FVS-421), cells were incubated with anti-CD32 antibody (BD Biosciences, Cat# 550270) for Fc receptor blocking (anti-CD32). Cells were stained with fluorochrome-conjugated antibodies (anti-rat antibodies: CD3 (eBioG4.18, G4.18) and CD4 (OX-38)) for surface marker detection. Cells were then fixed, permeabilized, and stained for intracellular FoxP3 (FJK-16s), Ki67 (SolA15), TNF-α (TN3-19.12), and IL-17A (eBio17B7). Samples were acquired on a BD LSRFortessa™ X-20 flow cytometer, and data were analyzed using FCS Express software (version 7.30.0030).

### Statistical Analysis

Data are expressed as Mean ± SD for each analysis, and statistical analyses using GraphPad Prism v11.1. Differences among groups (Naïve-EV, Hypoxia-EV, LPS-EV) were analyzed using one-way ANOVA followed by Sidak’s post hoc test. For cell viability and scratch assays, a two-way ANOVA followed by Dunnett’s test was conducted. A one-way ANOVA followed by Sidak’s post hoc test was used for T-cell experiments. The *α*-level for statistical significance was set at *p* < 0.05 for all statistical tests.

## Result

### Characterization of Human Placental Stem Cell-Derived Extracellular Vesicles

Extracellular vesicles derived from hPSC under different culture conditions (naïve, hypoxia-conditioned, and LPS-conditioned) were characterized to determine whether preconditioning influenced their physical properties. Nanoparticle tracking analysis (NTA) showed no statistically significant difference in EV concentration among the three groups. EV concentrations were 3.4 × 10^10^ ± 2.0 × 10^9^, 3.7 × 10^10^ ± 2.8 × 10^9^, and 3.7 × 10^10^ ± 4.7 × 10^9^ particles/mL for naïve-EVs, hypoxia-EVs, and LPS-EVs, respectively (*p* = 0.8195; **Fig. 1C**). Seeded hPSC numbers were confirmed to be similar between all groups and collections. Similarly, no statistically significant differences were observed in mean particle diameter (150.4 ± 1.0, 145.9 ± 2.5, and 149.9 ± 2.8 nm, respectively; *p* = 0.3239; **Fig. 1D)** or modal diameter (101.8 ± 4.8, 106.5 ± 7.3, and 106.7 ± 4.9 nm, respectively; *p* = 0.7930; **Fig. 1E**).

**Figure 1.**
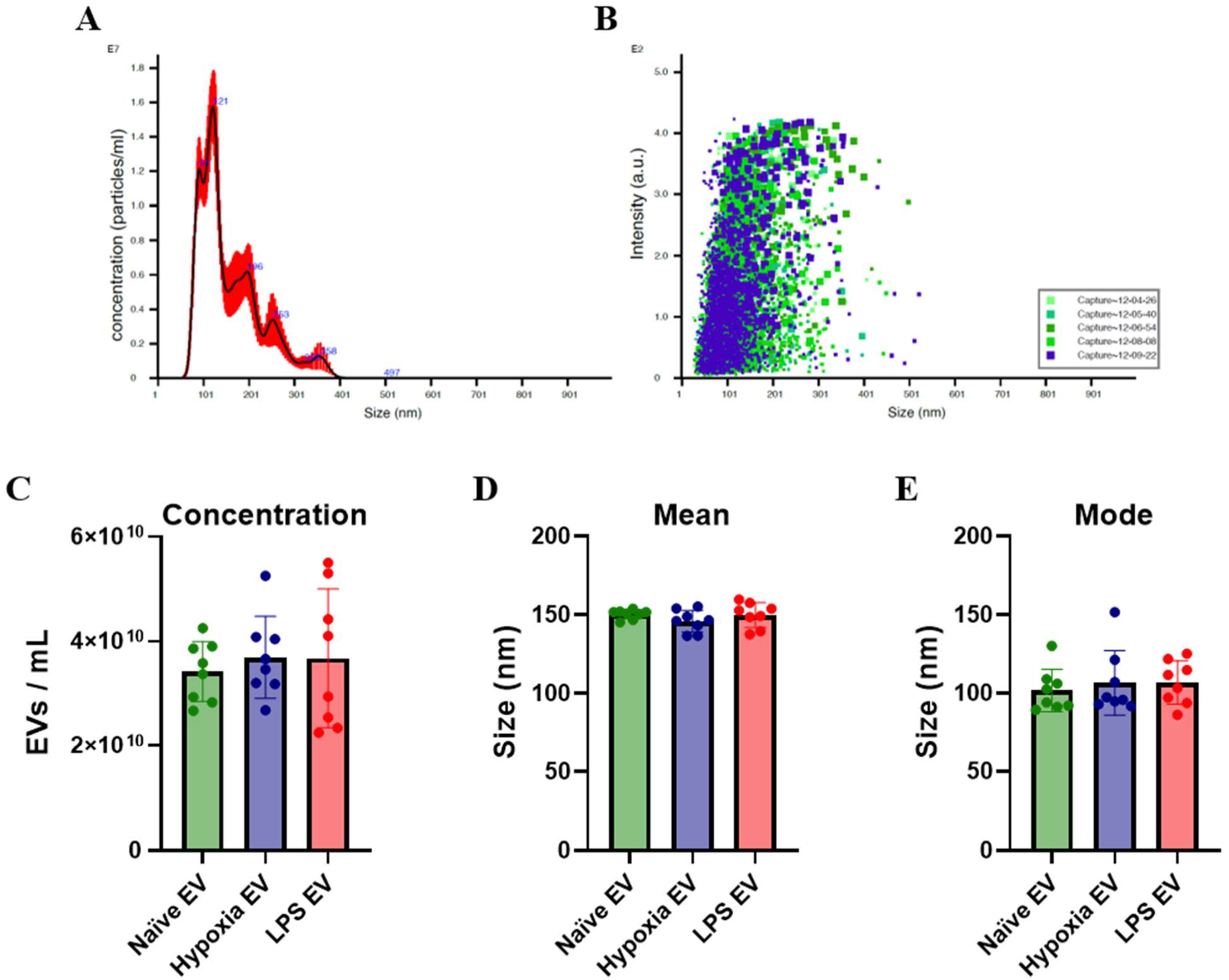
Nanoparticle tracking analysis of hPSC-derived EVs. hPSCs EVs isolated from naïve, hypoxia-conditioned, and LPS-conditioned hPSCs were characterized using NTA. (A) Representative particle size distribution and concentration profile. (B) Representative particle size and light-scattering intensity distribution. Quantitative comparisons of (C) EVs concentration (particles/mL), (D) mean particle diameter, and (E) mode particle diameter. No significant differences were observed in particle concentration, mean and mode diameter among the groups. n=8.

Transmission electron microscopy (TEM) following negative staining revealed hallmark features of extracellular vesicles; structures with rounded, cup-shaped, and dimple-like appearances across all three isolation conditions (**Fig. 2A–D**). These morphological features were consistent across conditions. Super-resolution imaging using the ONI Nanoimager further demonstrated the presence of the EV-associated tetraspanins CD9, CD63, and CD81 across all three groups (**Fig. 3A–J**). Analysis of individual particles revealed heterogeneity in tetraspanin expression (Fig. 3A-D), with approximately 24% EVs expressed all three transmembrane markers (CD9^+^CD63^+^CD81^+^). Approximately 52% expressed combinations of two markers (CD9^+^CD81^+^, CD9^+^CD63^+^, or CD63^+^CD81^+^), while the remaining 24% were positive for a single marker (CD9^+^, CD63^+^, or CD81^+^). Similar trends of no changes between EV groups were observed when analyzed according to their tetraspanin expression profiles and corresponding particle sizes (**Fig. 3E–G**). However, these observations were descriptive, as statistical comparisons were not performed due to the limited sample size (n = 2, with one measurement representing four pooled EV isolations used in the biological activity assays).

**Figure 2.**
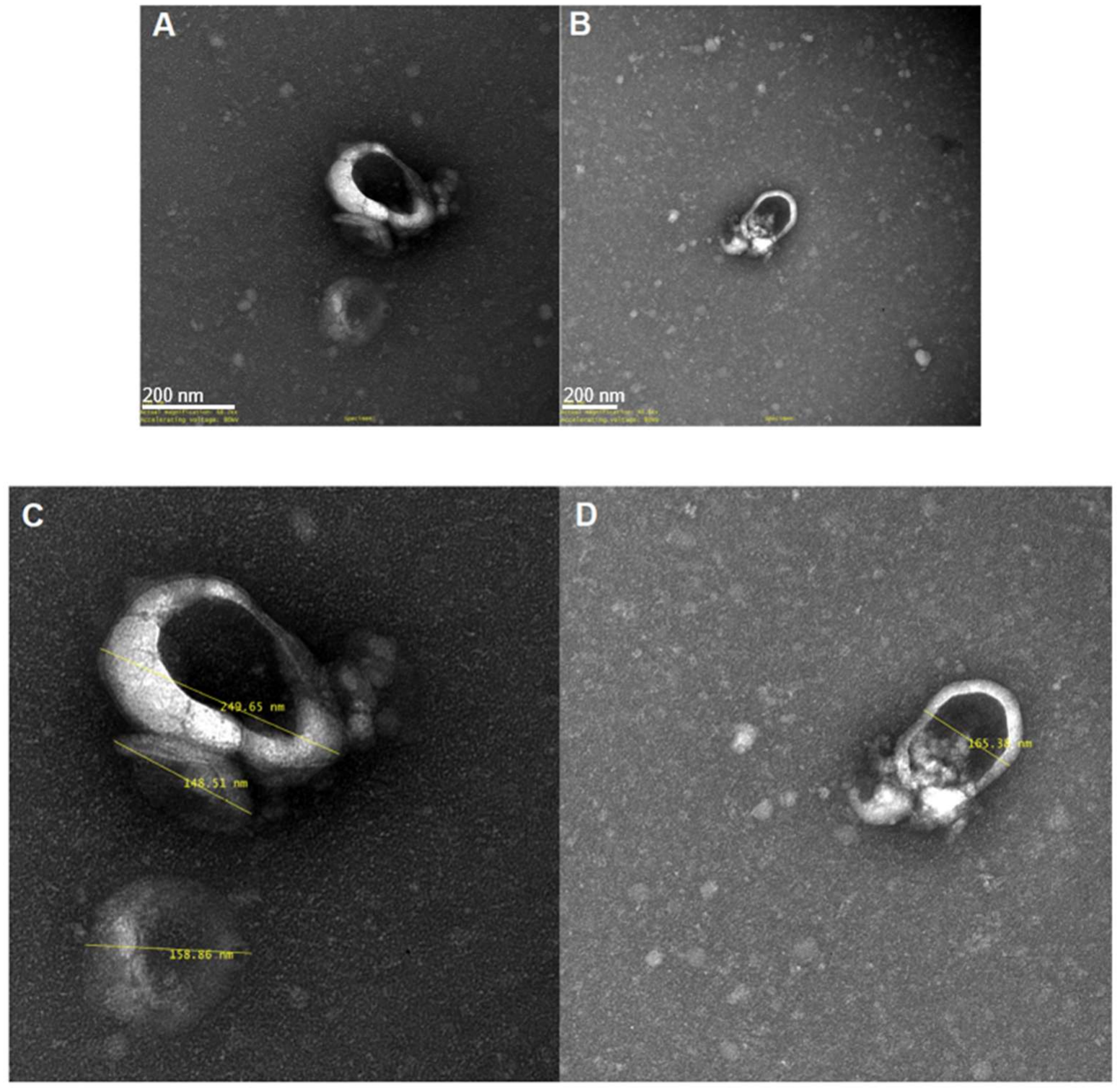
Transmission electron microscopy characterization of hPSC derived EVs. Representative TEM images of negatively stained EVs showing rounded, cup-shaped, and dimple-like morphology. Scale bar = 200 nm. **(A, B)** Representative images of EV morphology and **(C, D)** corresponding magnified images with particle size measurements (nm).

**Figure 3.**
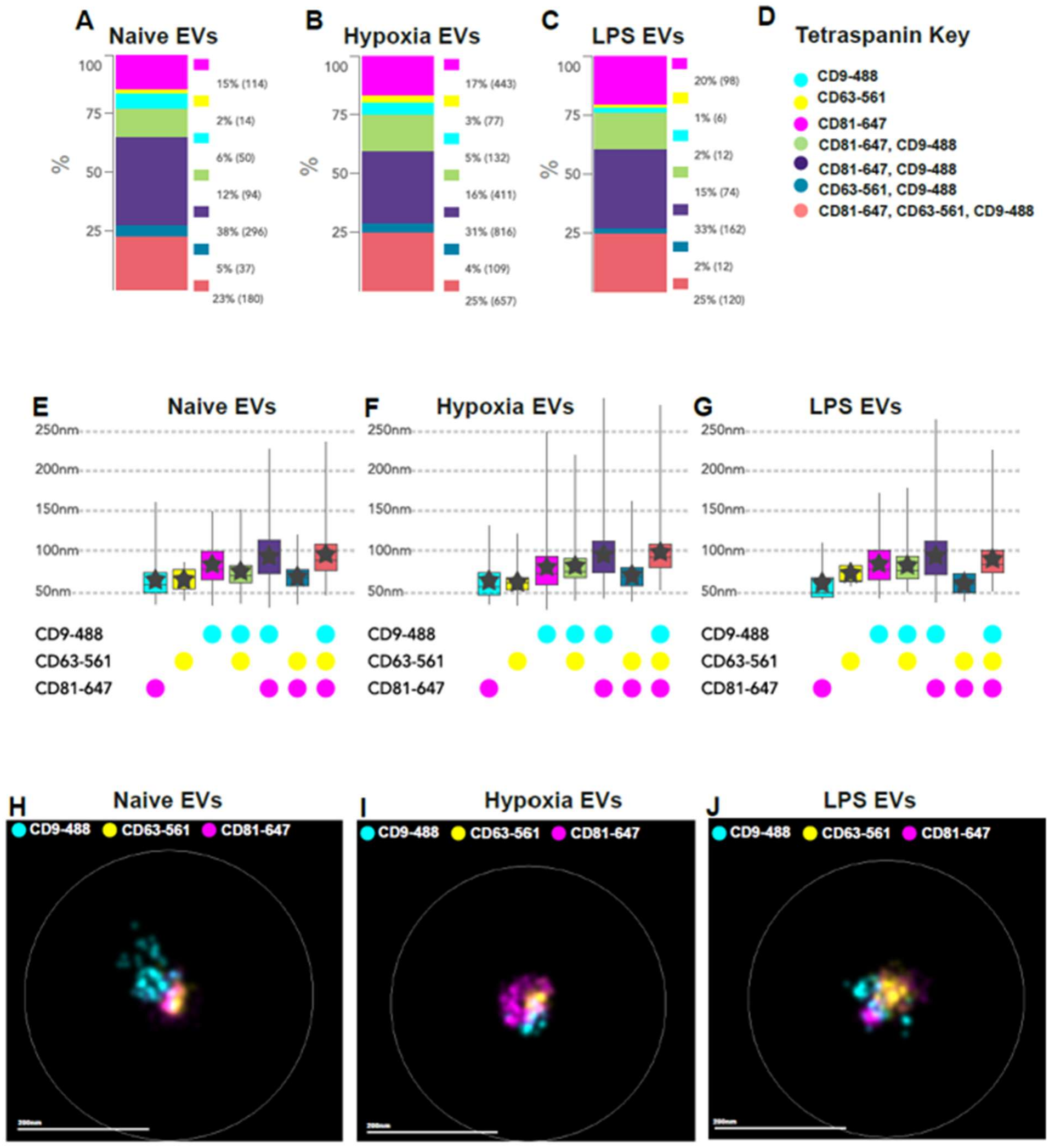
Tetraspanin expression and size distribution of hPSC derived EVs. hPSC Evs derived from naïve, hypoxia-conditioned, and LPS-conditioned were analyzed using super-resolution microscopy. Distribution of EV populations expressing single, double, or triple combinations of the tetraspanin markers CD9, CD63, and CD81 in naïve **(A)**, hypoxia-conditioned **(B)**, and LPS-conditioned **(C)**. Particle size distributions according to tetraspanin expression profiles across the three EV groups **(E–G)**. Representative super-resolution images **(H– J)** showing CD9 (cyan), CD63 (yellow), and CD81 (magenta) labeling in naïve **(H)**, hypoxia-conditioned **(I)**, and LPS-conditioned **(J)** EVs. Similar patterns of tetraspanin expression and particle size distribution were observed across the EV preparations. n = 2; Scale bar = 200 nm.

### Effects of hPSC-Derived EVs on Intestinal Epithelial Cell Viability, Wound Closure, and Nitrite Production

The effects of naïve and preconditioned EVs on intestinal epithelial cells were evaluated using cell viability, scratch wound healing, and nitrite production assays in a human fetal intestinal cell line (HIEC-6). The effects of naïve and preconditioned EVs on HIEC-6 cell viability were evaluated using the MTT assay. HIEC-6 cells exposed to increasing concentrations of LPS (0–10 µg/mL) showed a dose-dependent reduction in cell viability after 24 hours. HIEC-6 cells treated with naïve, hypoxia-conditioned, or LPS-conditioned EVs showed higher cell viability across all LPS concentrations compared with cells exposed to LPS alone. At each LPS concentration (2, 4, 6, 8, and 10 µg/mL), all three EV-treated groups showed significantly greater viability compared with the corresponding LPS-only control (*p* < 0.0001 for all comparisons; **Fig. 4A**). This protective effect was maintained even at the highest LPS concentration (10 µg/mL), where LPS alone caused the greatest reduction in cell viability.

**Figure 4.**
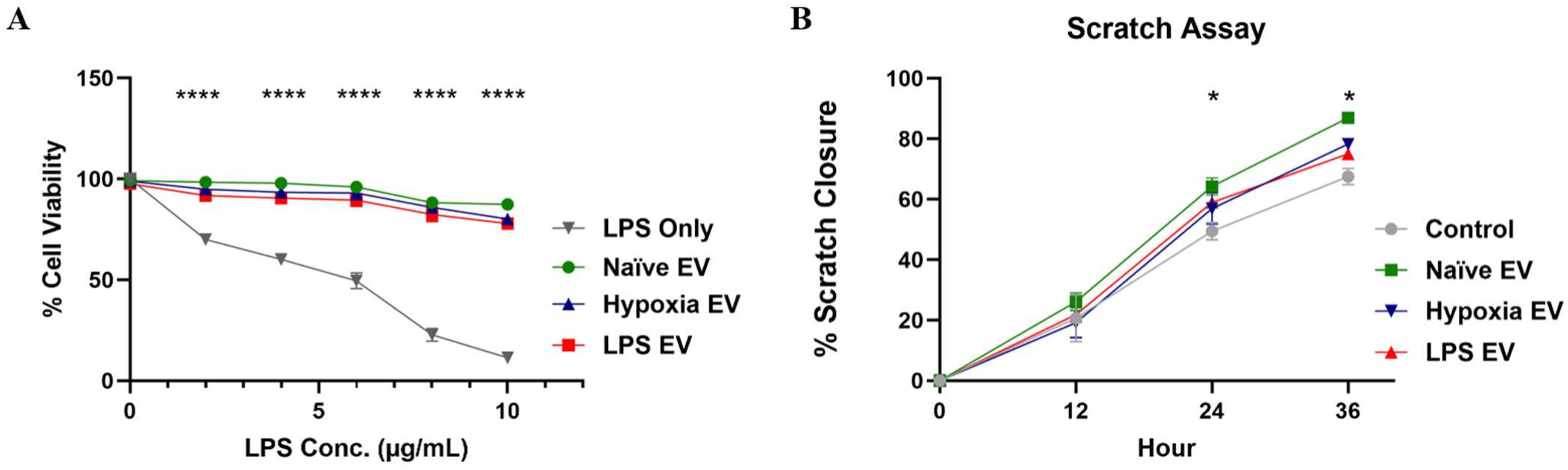
Effects of EVs Treatment on HIEC-6 cell viability and wound closure. **(A)** HIEC-6 cells were treated with increasing concentrations of lipopolysaccharide (LPS; 0, 2, 4, 6, 8, and 10 µg/mL) in the presence or absence of naïve EVs and preconditioned EVs. Cell viability was evaluated using the MTT assay, and results were expressed as a percentage of the untreated control. All EV-treated groups showed significantly higher cell viability compared with the corresponding LPS-only controls at each LPS concentration; ****p < 0.0001. **(B)** Scratch wound closure was monitored at 0, 12, 24, and 36 hours following EV treatment. All three EV groups showed significantly greater wound closure than untreated controls at 24 and 36 hours (\**p* < 0.05), n = 3.

To examine the effects of hPSC-derived EVs on epithelial wound closure, HIEC-6 cells were treated with naïve, LPS-preconditioned, or hypoxia-preconditioned EVs and wound closure was monitored at 0, 12, 24, and 36 hours. As shown in **Fig. 4B**, the naïve and preconditioned EVs-treated groups showed significantly greater wound closure than untreated cells at 24 hours (naïve-EVs, *p* < 0.0001; LPS-EVs, *p* = 0.0020; hypoxia-EVs, *p* = 0.0160). Wound closure continued to remain significantly greater in cells treated with naïve-EVs (*p* < 0.0001), LPS-EVs (*p* = 0.0162), and hypoxia-EVs (*p* = 0.0006) compared with untreated controls at 36 hours.

The effects of EV treatment on inflammatory responses within the intestinal epithelium were further examined by measuring nitrite accumulation in LPS-stimulated HIEC-6 cells using the Griess assay. Cells exposed to LPS (2 or 4 µg/mL) were treated with naïve, hypoxia-conditioned, or LPS-conditioned EVs. At both LPS concentrations, treatment with naïve, hypoxia-conditioned, or LPS-conditioned EVs significantly reduced nitrite levels compared with the corresponding LPS-only controls (*p* < 0.0001; **Fig. 5**). These findings indicate that naïve and preconditioned hPSC-EVs similarly reduced LPS-induced nitrite accumulation in intestinal epithelial cells. These findings demonstrate that hPSC derived naïve and preconditioned EVs were associated with improved epithelial cell viability and wound closure, along with reduced nitrite accumulation under LPS-stimulated conditions.

**Figure 5.**
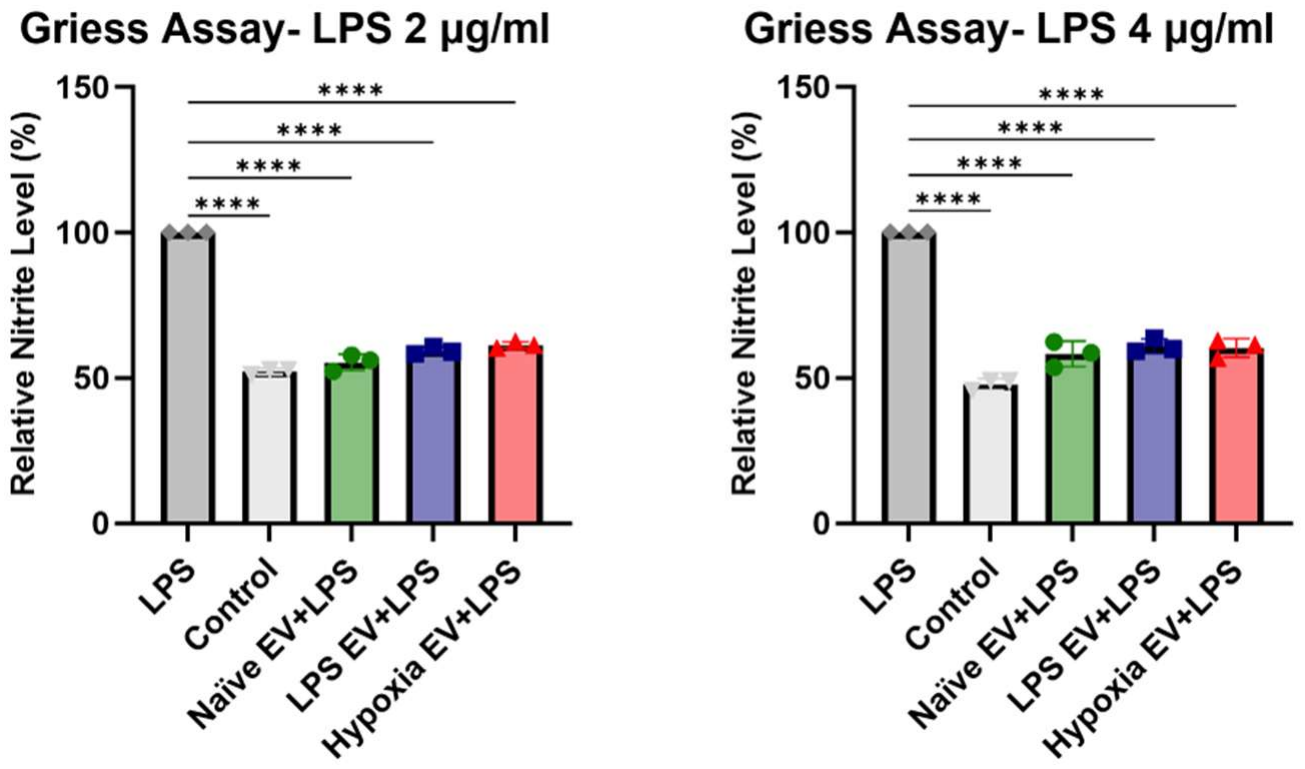
Effects of hPSC derived EVs on LPS-induced nitrite production in HIEC-6 cells. HIEC-6 cells were treated with LPS (2 or 4 µg/mL) in the presence or absence of naïve and preconditioned EVs. Untreated cells served as controls (light gray column= no LPS, no EVs). Nitrite levels were measured using the Griess assay and expressed relative to the corresponding LPS-only group (dark gray column= LPS, no EVs). \*\*\*\**p* < 0.0001.

### Effects of hPSC-Derived EVs on Neonatal Rat CD4+ T-Cell Responses

To assess the immunomodulatory effects of hPSC-derived EVs, primary neonatal rat splenic lymphocyte cultures were treated with naïve, hypoxia-conditioned, or LPS-conditioned EVs. Flow cytometric analysis showed no significant differences in the frequency of CD3^+^CD4^+^ T-cells among the three EV treatment groups. The frequency of TNFα-positive cells within the CD3^+^CD4^+^ T-cell population revealed no significant differences between naïve-EV and hypoxia-EV treatments (15.05 ± 5.77% vs. 13.76 ± 5.92%). However, a higher trend was observed following LPS-EV treatment (19.77 ± 6.44%), although the difference compared with naïve-EVs was not statistically significant. No significant difference was observed between hypoxia-EV and LPS-EV groups (**Fig.6A**). Similarly, the frequency of IL-17A-positive cells within the CD3^+^CD4^+^ T-cell population did not differ significantly among naïve-EV, hypoxia-EV, and LPS-EV treatments (3.4 ± 1.5%, 2.0 ± 1.0%, and 2.7 ± 0.4%, respectively; **Fig. 6B**). Mean fluorescence intensity (MFI) analysis also showed no significant differences in TNFα expression among naïve-, hypoxia- and LPS-EV groups (1282 ± 42, 1607 ± 269, and 1355 ± 29 AU, respectively), respectively; **Fig. 6D**). Likewise, IL-17A MFI remained comparable across treatments (4202 ± 1628, 3002 ± 458, and 2195 ± 5341 AU, respectively; **Fig. 6E**).

**Figure 6.**
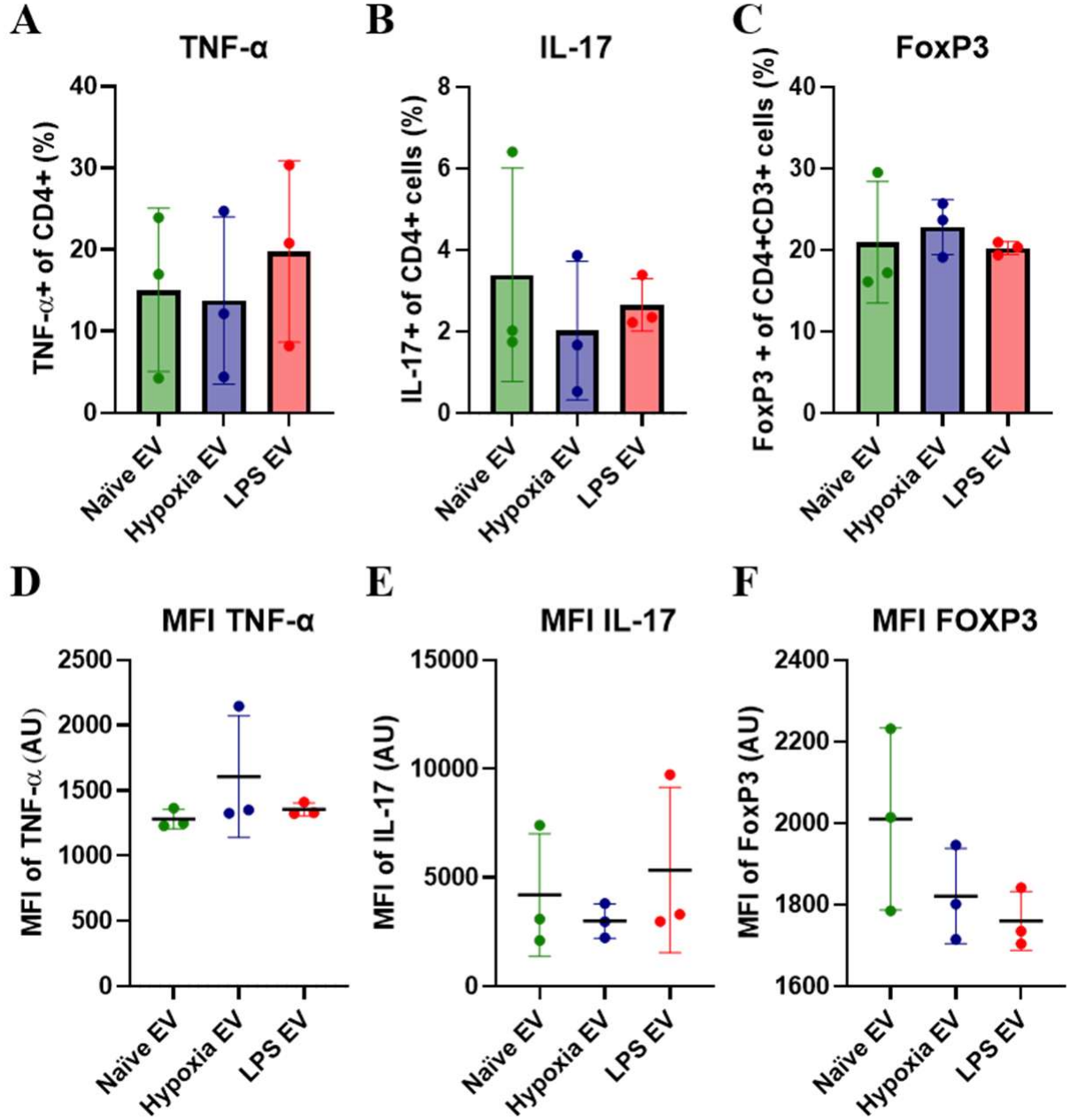
Effects of hPSC derived EVs on neonatal rat CD4^+^ T-cell responses. Neonatal rat splenic CD4^+^ T-cells were cultured with EVs derived from naïve, hypoxia-conditioned, or LPS-conditioned hPSC for 48 hours. Flow cytometric analysis was performed to assess the frequency of **(A)** TNF-α^+^, **(B)** IL-17A^+^, and **(C)** FoxP3^+^ cells within the CD3^+^CD4^+^ T-cell population. Mean fluorescence intensity (MFI) was measured for **(D)** TNF-α, **(E)** IL-17A, and **(F)** FoxP3. n = 3.

To further evaluate regulatory T-cell-associated responses, FoxP3 expression was examined within the CD3^+^CD4^+^ population. The frequency of FoxP3-positive cells was similar among naïve and preconditioned EV treatments (21 ± 4%, 23 ± 2%, and 20 ± 0%, respectively; **Fig. 6C**). FoxP3 MFI also showed no significant differences across the groups (2010 ± 129, 1821 ± 67, and 1760 ± 42 AU, respectively; **Fig. 6F**). Overall, these findings indicate that no significant differences in CD4^+^ T-cell cytokine or FoxP3 expression were observed among naïve and preconditioned EVs treatment groups.

## Discussion

The development of extracellular vesicle-based therapies for necrotizing enterocolitis (NEC) requires a better understanding of how EV production conditions influence their molecular composition and biological activity. In the present study, we investigated whether hypoxic or inflammatory preconditioning of human placental stem cells (hPSCs) could modify EV properties and potentially enhance their regenerative and immunomodulatory effects. Preconditioning did not significantly alter EV concentration, size distribution, or morphology among naïve, hypoxia-conditioned, and LPS-conditioned EVs. These data show that preconditioning did not significantly impact the EV formation and secretion dynamics as their physical characteristics remained unchanged. Despite molecular differences, all three EV preparations demonstrated beneficial and similar effects on intestinal epithelial cell responses. Further, their effects on the CD4^+^ T-cell markers examined remained similar with no significant differences between preconditioned EVs and naïve EVs. These findings suggest that while preconditioning influences the molecular composition of hPSC-derived EVs, the basic functional consequences of these changes were similar in these simplified *in vitro* studies. Furthermore, the full functional impacts of preconditioning require more comprehensive model systems of either more complex *in vitro* models or *in vivo* models and thus disease-specific changes between EV effects remain to be established.

An important consideration in developing EV-based therapeutics is the ability to generate preparations with consistent physical characteristics. EV size can influence biodistribution, tissue uptake, and clearance, making particle characterization an essential aspect of therapeutic development^9, 14^. In this study, ultrafiltration followed by size-exclusion chromatography was used to generate small EV-enriched preparations. Nanoparticle tracking analysis demonstrated comparable particle concentrations and size distributions across the three culture conditions, with mean particle diameters of approximately 146–150 nm. Transmission electron microscopy further confirmed the presence of characteristic vesicular structures, while super-resolution microscopy demonstrated the expression of CD9, CD63, and CD81 across the EV preparations. Similar patterns of tetraspanin heterogeneity were observed among the groups. Together, these findings indicate that hypoxic and LPS preconditioning did not produce detectable differences in the measured physical properties of the EV preparations, providing a consistent basis for comparing their molecular composition and biological activity.

All three EV preparations showed beneficial effects on HIEC-6 cells. Naïve, hypoxia-conditioned, and LPS-conditioned EVs maintained higher cellular metabolic activity following LPS exposure, increased scratch wound closure, and reduced nitrite accumulation compared with their respective controls. Similarly, primary healthy neonatal rat CD4^+^ T-cell cultures similar effects with no significant differences in TNFα, IL-17A, or FoxP3 expression among the three EV treatments. Although TNFα-positive T-cell frequency was numerically higher following LPS-EV treatment, this difference was not statistically significant. Previous studies have shown that EVs from LPS-stimulated cells can influence inflammatory responses^15^, suggesting that their effects may depend on the recipient-cell environment. Overall, neither preconditioning condition demonstrated a clear advantage over naïve EVs in the measured outcomes in these basic biological function studies in the cell culture models. Rather, these data show a consistent baseline of EV effects regardless of cell culture conditions. This suggests that more dynamic effects of preconditioning on EV impacts may require more complex model systems with interactions between the damages cell types such as enteroid and T-cell co-cultures or the *in vivo* NEC model.

Previous studies have reported improved therapeutic activity following stem-cell preconditioning, including enhanced intestinal protection with thrombin-preconditioned EVs in experimental NEC^4^ and altered regenerative and immunomodulatory activity following inflammatory priming. In our study, hypoxic and LPS preconditioning produced distinct protein profiles while maintaining a baseline impact in our selected measurements of functional responses. The relatively small sample sizes and limited simple *in vitro* models may have restricted our ability to detect differences between EV preparations. Furthermore, the biological contributions of individual proteins identified through proteomic analysis remain to be established. Future studies using more physiologically relevant NEC models and targeted protein analyses are needed to better understand the effects of preconditioning^1, 9^. Overall, hypoxic and LPS preconditioning altered hPSC-EV protein composition without significantly changing their physical characteristics. All three EV preparations showed beneficial epithelial responses and maintained effects on CD4+ T-cell expression. Taken together, these findings suggest that preconditioning influences EV molecular cargo, providing a basis for further investigation of their potential therapeutic applications in NEC.

## Contributions

Study conception and design: A.G., M.A.E. and V.W.; Data acquisition: A.G., M.A.E., Data analysis and interpretation: A.G., M.A.E and V.W.; Writing and editing the manuscript: all authors.

